# Open-source instrumented object to study dexterous object manipulation

**DOI:** 10.1101/2023.10.20.563288

**Authors:** David Córdova Bulens, Sophie du Bois de Dunilac, Benoit Delhaye, Philippe Lefèvre, Stephen J. Redmond

**Affiliations:** Biomedical Sensors & Signals Group, School of Electrical and Electronic Engineering, University College Dublin, Dublin, Ireland; Institute of Information and Communication Technologies, Electronics and Applied Mathematics (ICTEAM), Université catholique de Louvain, Louvain-la-Neuve, Belgium; Institute of Neuroscience (IoNS), Université catholique de Louvain, Brussels, Belgium

## Abstract

Humans use tactile feedback to perform skillful manipulation. When tactile sensory feedback is unavailable, for instance, if the fingers are anesthetized, dexterity is severely impaired. Imaging the deformation of the finger pad skin when in contact with a transparent plate provides information about the tactile feedback received by the central nervous system. Indeed, skin deformations are transduced into neural signals by the mechanoreceptors of the finger pad skin. Understanding how this feedback is used for active object manipulation would improve our understanding of human dexterity. In this paper, we present a new device for imaging the skin of the finger pad of one finger during manipulation performed with a precision grip. The device’s weight (300 g) makes it easy to use during unconstrained dexterous manipulation. Using this device, we reproduced the experiment performed in Delhaye et al. 2021a. We extracted the strains aligned with the object’s movement, i.e., the vertical strains in the ulnar and radial parts of the fingerpad, to see how correlated they were with the grip force (GF) adaptation. Interestingly, parts of our results differed from those in Delhaye et al. 2021a due to weight and inertia differences between the devices, with average GF across participants differing significantly. Our results highlight a large variability in the behavior of the skin across participants, with generally low correlations between strain and GF adjustments, suggesting that skin deformations are not the primary driver of GF adaptation in this manipulation scenario.

**Significance statement:** In this paper, we introduce a new device weighing 300 g and capable of imaging the skin of the finger pad of one finger during manipulation performed with a precision grip. This object is also capable of recording the forces and accelerations applied to the object. We reproduced the experiment performed in Delhaye et al. 2021a using this device. We extracted the strains aligned with the object’s movement to analyze how correlated these strains were with GF adaptation. The behavior of the skin across participants presented a large variability, and we observed low correlations between strain and GF adjustments in most participants. Our results suggest that skin deformations are not the primary driver of GF adaptation in this manipulation scenario.

## Introduction

We are capable of manipulating objects dexterously using our hands and fingers. To maintain a stable grasp, humans continuously adjust their grip force (GF), i.e., the force applied by the fingers normal to the contact surface, to counteract the load force (LF), i.e., the force caused by gravity and the inertia of the object tangential to the contact surface. GF adjustments are performed using a combination of predictions about the object’s response to the hand and arm movements and reactive control, driven by proprioceptive and tactile feedback Flanagan et al. 1993, White et al. 2008, Augurelle et al. 2003. Dependent on the movements being performed, the adjustments of GF can be either intermittently or totally synchronous with changes in LF (Grover et al. 2019, 2018). Moreover, tactile feedback can elicit a wide range of responses in the arm and hand muscles, affecting fast responses like short-latency reflexes and long-latency reflexes, or voluntary responses (Pruszynski et al. 2016, Crevecoeur et al. 2017).

Human dexterity is enabled by the number of degrees of freedom and the remarkable sensing capabilities of the hand. The human hand is innervated with several thousand mechanoreceptors that respond to skin deformations and provide information about the contact between the finger skin and the object being manipulated (Ackerley et al. 2014, Birznieks et al. 2001, Johansson and Birznieks 2004, Corniani and Saal 2020). It is assumed that the tactile information these mechanoreceptors provide plays a significant role in dexterous manipulation. Indeed, it has been shown that anesthetizing the tactile feedback at the finger pads significantly impacts dexterity (Augurelle et al. 2003, Monzée et al. 2003). However, what tactile feedback is actually received and how this feedback is used during object manipulation remains poorly understood.

The behavior of the finger pad skin when in contact with a flat glass plate has been extensively investigated in passive situations, i.e., when the finger is immobilized, and a glass plate is brought into contact with it. Indeed, imaging of the skin has shown that when the glass plate was displaced relative to the finger, the skin started slipping at the periphery of the contact, with the slip propagating inwards until all skin was slipping (André et al. 2011, Delhaye et al. 2014, Tada and Kanade 2004, Levesque and Hayward 2003). Analysis of the finger pad images allowed the measurement of how the different parts of the skin deform under these tangential and rotational loads (Delhaye et al. 2016, du Bois de Dunilac et al. 2022). These images highlighted patterns of compression and dilation at the edge of the propagating slip wavefront. Microneurography showed that skin mechanoreceptors fire in response to these evolving compressive/dilative strains and that the firing rate depended on the amplitude of the strain rate (Delhaye et al. 2021b). These results highlight that skin deformations occurring at the contact interface with an object are translated into sensory signals and provide feedback about the slip wavefront propagating across the finger pad skin. This information is potentially important in enabling the nervous system to control the grip force during active object manipulation to prevent slip.

Recently, studies have examined how the finger pad skin behaves when we actively manipulate an object (Delhaye et al. 2021a, Schiltz et al. 2021). These studies have highlighted the presence of strain patterns of compression and dilation appearing in specific regions of the finger. These strains could provide important information about the object being manipulated, the movement being performed, and the contact state. However, the device used in these studies and presented in Delhaye et al. 2021a has two important limitations stemming from the large weight of the device (500 g), preventing it from being manipulated easily in a precision grip and requiring a counterweight to allow participants to manipulate it more easily. First, this counterweight creates a discrepancy between the device’s apparent weight and its mass/inertia. Second, the counterweighting cable constrains the device to vertical movements, limiting the experiments that can be performed with it to involve strictly vertical motion.

In this paper, we present a new open-source device capable of imaging the skin of the finger pad of one finger during manipulation performed with a precision grip; we name the device InOb, for instrumented object, in the rest of the manuscript for brevity. InOb weighs only 300 g and is therefore easily manipulated by human participants. Using InOb, we performed an experiment where participants performed vertical oscillations, similar to the experiment performed in Delhaye et al. 2021a. We observed a large variability in the behavior of participants’ skin during the task, with strain levels differing significantly. Across all participants, we found no clear relationship between skin strains and grip force adjustments, suggesting that strains might not be the main driver for GF adjustments in this periodic task. Indeed, the periodicity of the task likely allows the central nervous system to predict the changes in LF and adjust GF synchronously.

## Methods

### Hardware and design

The InOb was designed to image the skin of one finger pad during active manipulation. For this reason, InOb is dimensioned to be manipulated in a precision grip; i.e., pinched between the thumb and the index finger. Its optical system allows the imaging of the contact that one finger pad makes with the transparent plate. A direct illumination system was selected as it results in high contrast between fingerprint ridges and valleys and leads to a compact structure (Bochereau et al. 2017). The custom optical system (Fig. 1C) is composed of a light source (Neewer SL-12), a half mirror (Edmund Optics part #43-359), a camera (Raspberry Pi Camera v2), and a lens with adjustable focal length (CWBL2.812-3MP-C, 8-13.5mm lens). From the light source, half the light goes through the half-mirror, reflects off the transparent plate in contact with the finger, and goes back to the half-mirror, where half of the light coming from the plate is reflected into the camera lens. The internal walls of InOb were painted using Black 3.0 paint (Stuart Semple) to minimize light reflections from the internal walls. This technique has been used to image the finger of humans during static loading (Delhaye et al. 2014, 2021b, Bochereau et al. 2017, Willemet et al. 2022, Khamis et al. 2021) and dynamic manipulation (Delhaye et al. 2021a).

**Fig. 1.**
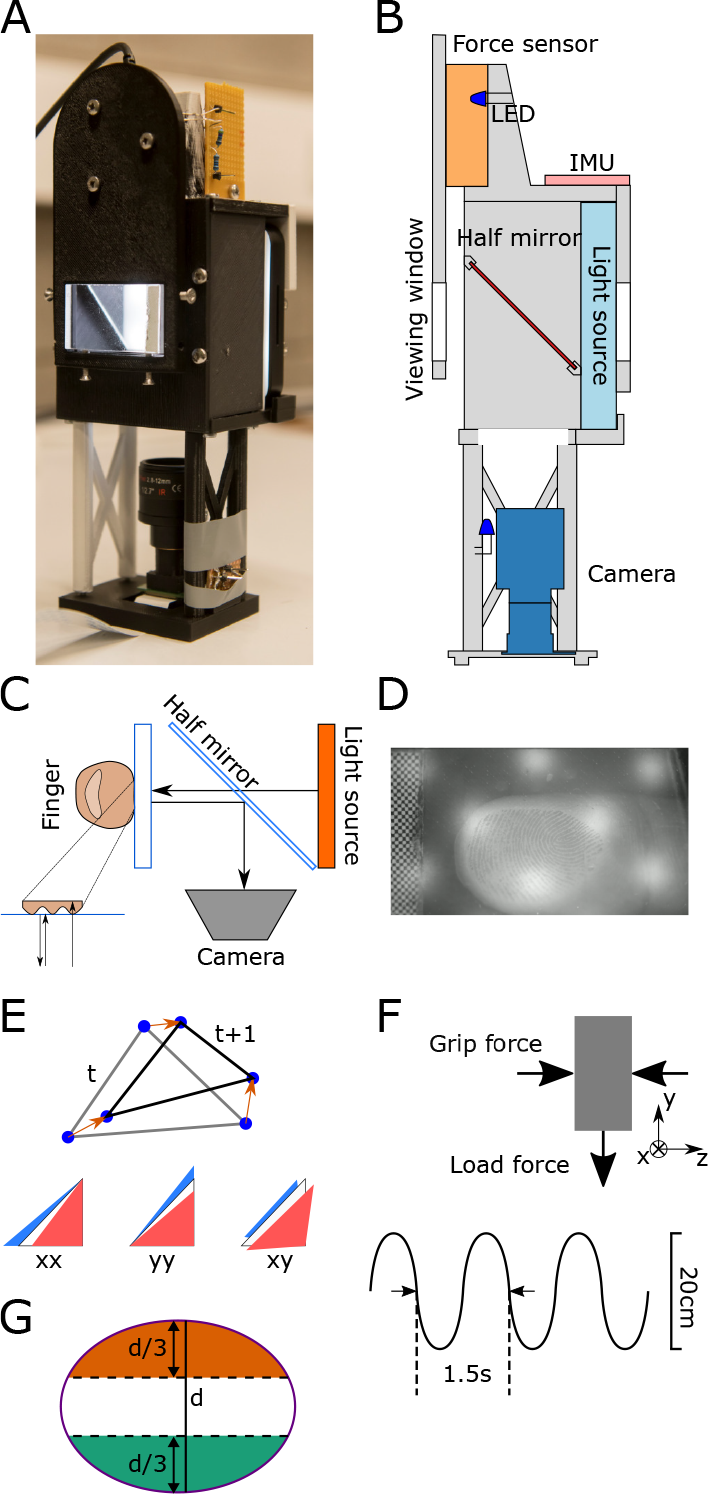
Methods. A) Photograph of the object showing the glass plate, connected to the force sensor, through which the finger is going to be imaged. B) Side view of the instrumented object. The light source, half mirror, viewing window and camera represent the optical system allowing for the finger in contact with the viewing window to be imaged. The force sensor measures the 3D forces and torques. The LEDs are used to verify the synchronization of the force signals and the videos. C) Imaging principle of the instrument object. A light source (light-blue rectangle) illuminates the finger-glass contact. The image of the finger is reflected into the camera by a half-mirror. D) Image of the finger captured by InOb. E) Triangle shape evolution from frame t to frame t + 1 and corresponding representation of compression (red triangle) and dilation (blue triangle). F) Schematic of the forces acting on the object as well as the movement frequency and amplitude. G) Schematic view of the contact area with the division in 3 parts for analysis of the vertical strains of the finger (*ϵ*_yy_). The green and orange regions represent the ulnar and radial parts of the finger, respectively.

Furthermore, InOb is equipped with a six-axis force/torque sensor (ATI Mini 40, ATI-IA, USA) connected to the plate in contact with the finger, measuring the forces and torques applied to the transparent plate during manipulation. A 9degrees of freedom (DOF) inertial measurement unit (IMU, BNO055) (Fig. 1A and 1B) measures the accelerations and orientation of the object. Finally, InOb takes advantage of the (undocumented) light sensitivity of ATI sensors to ensure that the force and video signals are synchronized; at the start of the trial two LEDs, one next to the ATI sensor and one placed next to the camera lens, are flashed simultaneously for half a second providing a synchronous time event in the video and the force data recordings. The flashing LED leads to a change in the force along the x-axis of 0.05N. All other parts of the device are 3D printed in Tough PLA using a Makerbot Method X printer. The total weight of the object is 300 g and the dimensions are 75*×*77 *×* 225 mm (width*×* length *×* height).

The images from the camera are captured with a resolution of 720 *×* 1280 px (30 px/mm in the scene) at 60 fps. A lens with an adjustable focal length of 8-13 mm was chosen to achieve a focused image of the participant’s finger. Forces and torques are acquired at 200 Hz, and acceleration and orientation (Euler angles) are acquired at 100 Hz. The data is gathered using a Raspberry Pi 4B and Python 3.7. The acquisition code is provided in the following link.

### Pilot study

#### Participants

Ten participants (4 female, age 30.3 *±* 4.38 years) participated in this experiment. The experimental protocol was approved by the ethics committee of University College Dublin and informed consent was obtained from all participants prior to commencement.

#### Experimental protocol

To test whether InOb was capable of accurately imaging the finger pad skin of the index finger, while synchronously acquiring the forces and accelerations applied on the object during a manipulation task, we performed an experiment where participants lifted the object in a precision grip, with the index finger touching the viewing window, and performed vertical oscillations of 20 cm peakto-peak amplitude at a rate of 0.75 Hz for 20 s (Fig. 1F). Participants were instructed to reach the top and bottom of the movement at each beep of a metronome, which helped regulate the frequency of the movement. The amplitude of the movement was visually indicated using two visual markers placed adjacent to the trajectory of InOb. This experiment followed the same protocol as Delhaye et al. except that the weight was not compensated. After each trial, participants rested for 30 seconds before the start of the next trial. During this time, the viewing window was cleaned. In total, each participant performed 15 trials. The viewing plate was cleaned between trials.

After the 15 trials, the coefficient of friction (*μ*) of the contact between each participant’s finger and the viewing window of InOb was evaluated using the procedure described in Barrea et al. (2016). This procedure was conducted using InOb.

### Data analysis

#### Force, torque and acceleration analysis

Force, torque, acceleration, and orientation data were band-pass filtered using 4th-order low-pass Butterworth filters with cut-off frequencies of 40 Hz using zero-phase filtering. GF was estimated as the z-axis force measurement of the ATI sensor, and LF was estimated as twice the y-axis force of the ATI sensor (Fig. 1F). The start of a trial, i.e., the grasping of the object, was identified using the first peak in GF, and the end of a trial, i.e., the release of the grasp, was identified by the last peak in GF. The vertical velocity of the object was estimated by numerically integrating the z-axis acceleration measured by the IMU in the object’s reference frame. Individual oscillations (periods) were separated using the peaks of the vertical velocity.

We extracted the average GF and LF for each oscillation in order to analyze whether participants adapted their GF within and across trials. Using the coefficient of friction, we computed the friction limit as *μ*^*−*1^ and then computed the slip force (SF), i.e., the minimum GF required to maintain the object in a stable grasp for a given LF, as SF = LF *μ*^*−*1^. SF allowed us to compute the excess force applied by participants during the trials; i.e., the safety margin (GF *−* SF). We also extracted the GF modulation (ΔGF) across oscillations as the difference between the maximum and minimum GF in a single oscillation (max(GF) *−* min(GF)).

#### Image analysis

To improve the contrast of the fingerprints on the video, each frame was spatially filtered using a 2D Gaussian bandpass filter (0.4 mm). Features in the contact area were detected and tracked across frames using the LucasKanade-Tomasi algorithm (Lucas and Kanade 1981, Shi and Tomasi 1994, yves Bouguet 2000). The distribution of the tracked features was uniform, with the size of the triangles being 0.123 *±* 0.012 mm^2^ (mean*±* SD). These features were used to compute the skin displacement field. Using these features, we tessellated the contact area into triangular skin elements by performing a Delaunay triangulation. The contact area was segmented using a semi-automatic machinelearning algorithm.

The size of the contact area was determined by counting the pixels inside the segmented region and converting the result to mm^2^ using the conversion ratio of 30 px/mm given by the checkerboard pattern. This contact area can be modified by skin breaking or making contact, and skin deformations. To estimate the amount of skin lifting off or coming into contact at the edge of the contact area, we summed the areas of all the triangles of skin entering or leaving contact. A triangle was considered inside the contact area if its three vertices are located inside it. We computed the surface deformations leading to area changes by computing the area change of all Delaunay triangles in the contact area from one frame to the next.

The amount of skin slipping in each frame of the video was measured by considering that a triangular skin element had slipped if its centroid had moved more than 0.5 pixels across five consecutive frames. The stick ratio (SR) was then computed as the ratio of the area of skin stuck, i.e., the skin that is in the contact area but has not been detected as slipping as a fraction of the total contact area. We labeled each oscillation as ‘slip’ if the stick ratio dropped below 0.5. We used this label to compare the amplitude of the GF change in each oscillation between oscillations labeled as slip and trials with no slip.

The deformations of each triangular skin element were quantified by a strain tensor, *ϵ*, having three independent values: the normal strain along the x-axis (*ϵ*_xx_), the normal strain along the y-axis (*ϵ*_yy_) and the shear strain (*ϵ*_xy_) (Fig. 1D). These strains (two-dimensional Green-Lagrange) were computed as the change in the shape of each triangular element between consecutive frames (Fig. 1E). The complete details of the strain computation can be found in Delhaye et al. 2016 and du Bois de Dunilac et al. 2022.

As the movement was vertical, and based on the strain patterns observed by Delhaye et al. 2021a, we divided the contact area into three segments in the ulnar to radial direction (Fig, 1G). Indeed, we expected strains to appear mainly in the radial and ulnar parts of the finger (Delhaye et al. 2021a). The division between these segments was performed by computing the longest axis in the ulnar to radial direction of the contact area and dividing that length equally into three parts. We then computed the average of *ϵ*_yy_ in each region, weighted by the size of each triangular skin element, named *ϵ*_radial_ and *ϵ*_ulnar_ for the radial and ulnar part of the finger, respectively. These weighted averages allow us to identify the patterns and the amplitude of the strain happening in the upper and lower parts of the finger pad. For each oscillation, we computed the amplitude of *ϵ*_ulnar_ and *ϵ*_radial_ to study how much skin deformation happened within each oscillation cycle.

#### Statistical analysis

To analyze how participants adapted their GF within and across trials, we computed a repeated measures ANOVA with the trial number (#_*trial*_) and oscillation number (#_*osc*_) as within-participant factors and the average GF for each oscillation as the independent variable. We also performed repeated measures ANOVA with the same dependent variables, but with the average LF for each oscillation as the independent variable, to test whether LF changes significantly across or within trials.

To analyze the effect of slips on GF, we ran a paired t-test with the ΔGF in each oscillation for trials labeled as ‘slip’ and those labeled as ‘no slip’. We expected to observe that oscillations labeled as ‘slip’ would present a greater ΔGF than the oscillations labeled as ‘no slip’.

We computed the amplitude of the change of the weighted average of the strain in both the radial and ulnar parts of the finger in each oscillation, thus getting a single value per oscillation. To do so, for each oscillation we computed the difference between the maximum and minimum of *ϵ*_radial_ for the radial part of the finger, and the difference between the maximum and minimum of *ϵ*_ulnar_ for the ulnar part of the finger. We computed the amplitude of the change in contact area (normalized by the initial contact area of each trial) caused by the peeling and laying of the skin in each oscillation and trial. We also computed the change in contact area caused by compression and dilation of the skin for each oscillation and trial. To do so, we computed the difference between the minimum and maximum change in contact area caused by peeling & laying and by compression & dilation for each oscillation (Δ_*P eel*_). We also computed the minimum stick ratio SR_min_ for each oscillation to analyze how compromised the grip is. We performed repeated measures ANOVA on all these parameters with trial and oscillation as within-participant factors.

To investigate whether strain patterns could lead to adjustments of GF and explain the continuous modulation we observed, we computed cross-covariance using a sliding time window to analyze how well-synchronized both signals were across a complete trial, and how the time lag between both signals evolved across trials. We used a time window of 2.25 s to perform the cross-covariance, and this window was moved by 0.1 s at each step. This window size was selected to include two oscillations and reduce the variability of the cross-covariance measure. Indeed, if certain strain patterns trigger adjustments in GF, then we would expect the strains to change before a GF adjustment with a maximum covariance value for a negative time delay, i.e., GF lags strain. We performed a similar analysis between GF and peeling/laying, GF and compression/dilation, and GF and LF to investigate the coupling between GF and these different variables. Indeed, as LF is the main cause of strains appearing, we assumed we would observe similar covariances between Strains & GF, and LF & GF.

To further explore changes in the relation between strains and GF, we extracted the correlation coefficient between GF and *ϵ*_radial_ and *ϵ*_ulnar_ for each oscillation of each trial of every participant. We then computed a repeated measures ANOVA with these correlations as the dependent variable, and #_*osc*_ and #_*trial*_ as the independent variable.

## Results

During a typical trial, participants lifted the object and performed vertical oscillations with an amplitude of 20 cm peakto-peak with a frequency of 0.75 Hz for 20 s for a typical amount of 15 oscillations. Measured accelerations were in the range of *±*4 m.s^*−*2^ (Fig. 2B). The changes in the acceleration applied on the object led to the load force changing accordingly during the movement (Fig. 2A), with LF varying between 2 and 4 N. During a trial, participants adjusted their grip force continuously to keep a safe grasp during the whole movement (Fig. 2A). The grip security is reflected in the ratio between GF and LF which remains above the friction limit 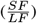 and remains stable across an entire trial (Fig. 2C), indicating that participants applied a large enough GF to prevent full slip. The stick ratio stayed close to 1 for most oscillations but presented some significant drops close to 0.5 for some oscillations (Fig. 2D), which suggests that participants did not allow large levels of slip to occur during the trial. Skin lost contact and came into contact with the glass plate as oscillations were performed (Fig. 2E). Peeling and laying accounted for changes of around 5% of the initial contact area across oscillations for most participants, and global compression and dilation also accounted for around 1% (Fig. 3B).

**Fig. 2.**
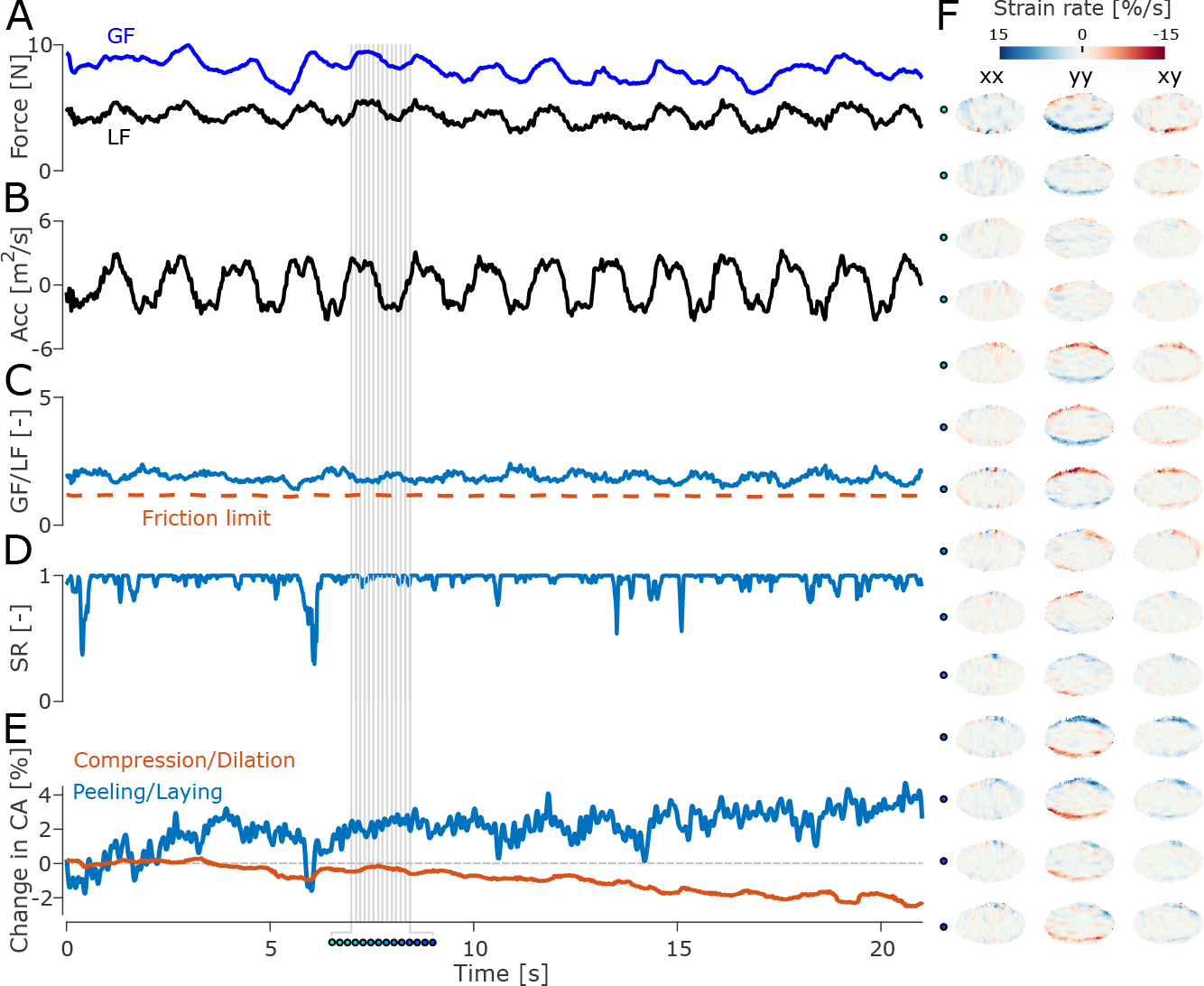
Forces, acceleration (Acc), stick ratio (SR), peeling, and strain rate for a single trial of an exemplar participant. A) Grip force applied (blue line) and load force experienced (black line) by the index finger of the participant during the trial. B) Vertical acceleration of the instrumented object measured by the IMU. C) Ratio of the grip force relative to the load force (black line) and friction limit (orange dashed line) which is estimated at the end of each experimental trial. D) Stick ratio of the skin; i.e., the region of the contact area where the skin is stuck to the plate as a proportion of the total contact area. E) Peeling and laying, and compression and dilation of the finger; i.e., amount of skin coming into and losing contact with the glass plate. F) Horizontal (*ϵ*_xx_), vertical (*ϵ*_yy_), and shear (*ϵ*_xy_) strain rates on the finger surface for a single oscillation. The filled-in dots represent the time stamps of the oscillation marked by vertical gray lines in panels A to E.

**Fig. 3.**
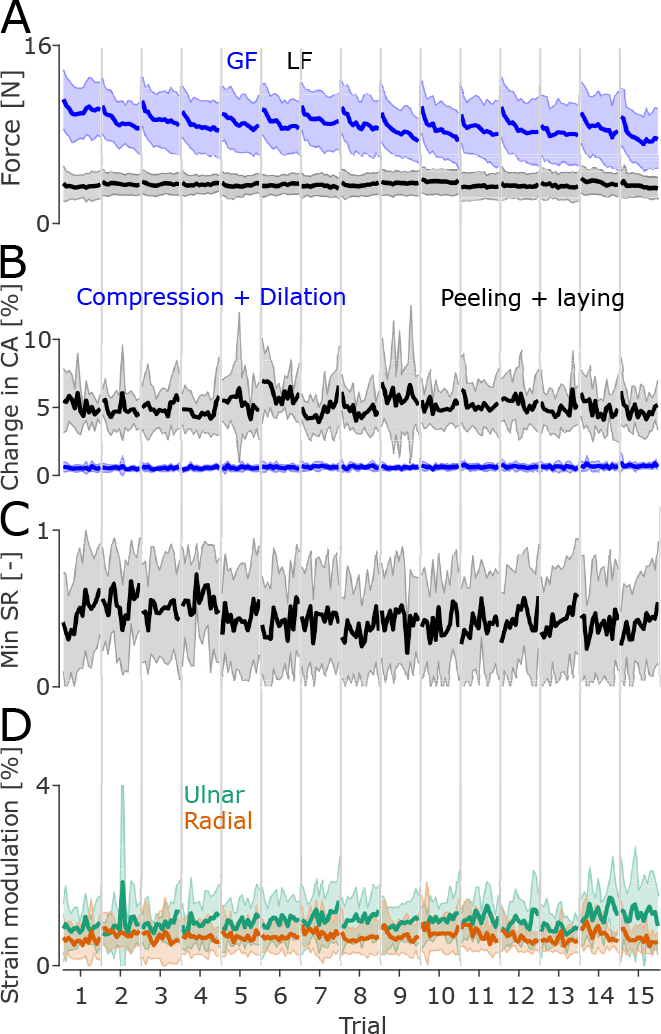
Mean GF and LF evolution, peeling amplitude, minimum slip ratio, and strain amplitude for each oscillation, aggregated across all trials. A) GF (blue line) and LF (black line) evolution within and across the 15 trials in the experiment aggregated across all participants. The shaded areas represent the standard deviation. B) Mean (black line) and SD (gray area) peeling amplitude across participants for each oscillation and each trial. C) Mean (black line) and SD (gray area) of the minimum slip ratio of each oscillation across participants. D) Mean (solid line) and SD (faded area) of the vertical strain modulation for the radial part of the finger (orange line) and the ulnar part of the finger (green line) of each oscillation across participants.

### Grip force and safety margin

The mean GF of each oscillation progressively decreased within a trial, often starting at forces of around 10 to 12 N before decreasing to forces of around 8 N by the last oscillation in a trial (Fig. 2A and 3A). Moreover, GF also progressively decreased across trials (Fig. 3A). Indeed, a repeated measures ANOVA showed that GF did vary significantly with #_*trial*_ (df = 14, F = 2.21, p = 0.024), and with #_*osc*_ (df = 13, F = 17.563, p < 0.001). There was no significant interaction (df = 182, F = 1.15, p = 0.11). As expected, the mean LF remained at similar levels across oscillations and trials. A repeated measures ANOVA run with LF as the independent variable showed no effect of #_*trial*_ (df = 13, F = 1.95, p = 0.054), #_*osc*_ (df = 14, F = 0.69, p = 0.76) and no interaction (df = 182, F = 0.833, p = 0.92). Therefore, as GF decreased within and across trials and LF did not, the amount of excess force exerted by participants through their grip decreased (Fig. 3A), suggesting that participants progressively adjusted their grip to the load exerted by the object, or fatigued during a and across trials.

Overall, all participants but one applied a GF that was, on average, well above (mean *±* SD excess force GF-SF of 4.61*±*2.58 N across participants) the slip force required to maintain a stable grasp, as shown by all participants GF staying above the dashed line in Fig. 4A. Indicating they all applied a substantial safety margin to ensure the object would not slip.

**Fig. 4.**
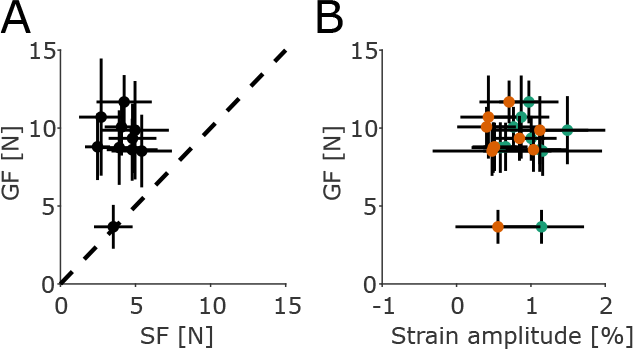
A) Participants’ mean GF and mean SF (black dots). The error bars represent the standard deviation. The dashed black line represents the slip force limit, below which slip will occur. B) Participants’ mean GF and mean strain amplitude for each oscillation for the ulnar (green dots) and radial (orange dots) parts of the finger.

### Peeling and compression of the skin

Skin regions lost contact with the plate and came into contact with the plate regularly during manipulation (Fig. 2E). Indeed, the amount of skin in contact with the plate varied by an average of 5% during trials. A repeated measure ANOVA computed on the amplitude of the change in contact area due to peeling/laying during an oscillation showed a significant effect of #_*osc*_ (df = 13, F = 2.11, p = 0.04), no significant effect of #_*trial*_ (df = 14, F = 0.75, p = 0.75), and a significant interaction (df = 182, F = 1.38, p = 0.003). This suggests that peeling/laying varied across oscillations within a trial (mean *±* SD in the first oscillation 0.0541 *±* 0.0212 and in the last oscillation 0.0468 *±*0.0259) but remained at similar levels across all trials (Fig. 3B). The significant interaction suggests, however, that peeling evolved with oscillations differently across trials, with some trials showing a decrease in the effect of peeling/laying on the contact area across oscillations and others showing an increase across oscillations.

Compression/dilation had a much smaller effect when compared with peeling/laying, accounting for changes in the contact area of about 1% (Fig. 3B). A repeated measures ANOVA computed on the amplitude of the change in contact area due to compression/dilation showed no significant effect of #_*osc*_ (df = 13, F = 0.69, p= 0.76), no significant effect of #_*trial*_ (df = 14, F = 1.3, p = 0.25) and no significant interaction (df = 182, F = 0.81, p = 0.95). These results suggest that changes in the contact area due to compression/dilation did not vary across oscillations or trials.

### Stick ratio

The stick ratio varied significantly across participants, with most of the skin remaining stuck for most of a trial with some momentary sudden drops in stick ratio (Fig. 2D) for most participants. It is worth noting that the levels of slip varied across participants, with the minimum stick ratio averaged across participants being close to 0.5 (Fig. 3D). A repeated measures ANOVA on the minimal SR observed on each oscillation did not show a significant effect of #_*osc*_ (df = 13, F = 0.53, p = 0.89), no significant effect of #_*trial*_ (df = 14, F = 1.69, p = 0.09), and no significant interaction (df = 182, F = 0.80, p = 0.96). This suggests that, despite participants reducing their grip force as the experiment progressed, the amount of slip they allowed during manipulation did not significantly increase as a consequence because they stayed above the friction limit throughout the experiment.

We labeled oscillations as ‘slip’ when the stick ratio dropped below 0.5 and as ‘no slip’ otherwise. We extracted the amplitude of the change in GF for each oscillation and compared it across these two labeled sets. Oscillations labeled as ‘slip’ showed a greater amplitude of change in GF compared to oscillations labeled ‘no slip’ (Fig. 5). A paired t-test showed a significant difference between these two conditions (p < 0.001, ci = [0.9085 - 1.0629]).

**Fig. 5.**
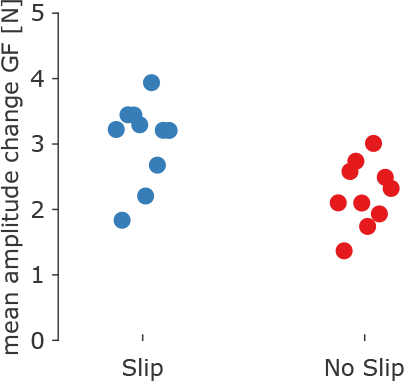
Mean difference in GF change amplitude for oscillations labeled as ‘slip’ (light blue dots) and ‘no slip’ (red dots) for each participant.

### Local strain patterns

During the course of a single oscillation, we observed repeating patterns of local compression and dilation happening on the skin. The vertical component of the strains (*ϵ*_yy_ in Fig. 2F) showed a repeating pattern of local compression and dilation in the radial and ulnar parts of the contact area. Indeed, we observed that the radial part of the contact area experienced skin compression when the acceleration was at its largest, and vertical skin dilation at the low point of the movement, and vice-versa for the ulnar part of the contact area, with this pattern repeating in each oscillation. Participants who exerted a larger GF did not show a smaller amplitude of strain in the ulnar and radial parts of the finger (Fig. 4B).

To better characterize this pattern, we computed the weighted average of the *ϵ*_yy_ in the top and bottom sections of the finger. We then detrended these averages and performed a numeric integral to obtain the average strain deformation in these two finger parts over time. A repeated measures ANOVA on the amplitude of *ϵ*_ulnar_ showed no significant effect of #_*trial*_ (df = 14, F = 1.08, p = 0.39) or #_*osc*_ (df = 13, F = 0.33, p = 0.98), and no interaction (df = 182, F = 0.8, p = 0.957). Similarly for *ϵ*_radial_, we found no significant effect of #_*trial*_ (df = 14, F = 0.577, p = 0.86) or #_*osc*_ (df = 13, F = 0.49, p = 0.91) and no significant interaction (df = 182, F = 1.032, p = 0.39). These results suggest that the strain amplitude did not vary across trials nor oscillations (Fig. 3C), even though GF reduces across oscillations and trials.

### Covariance between strains, peeling/laying, compression/dilation, LF and GF

We computed a time-windowed cross-covariance analysis for each trial to verify how the covariance between GF and strains evolved within a trial. For visualization purposes, we averaged these cross-covariance results across all trials for each participant. Moreover, we used the amplitude of the strains in the radial and ulnar parts of the finger as an indicator of the noise of the cross-covariance analysis (Fig. 6A). If strain patterns in the ulnar and radial parts of the finger happen on the skin at the same frequency as the movement, we would expect a high covariance between GF and these strains. Moreover, if strains and tactile feedback drive GF adjustments, we would expect the strains to lead GF time-wise leading to a covariance peak at a positive lag. We observed that the covariance between strains and GF varied widely across participants (Fig. 6B). Some participants presented a large covariance between strains and GF, while others showed small to no covariance (Table 1). The lag between GF and strains also differed greatly across participants, with a global average of -4.8*±*528 ms and -7.6 *±* 541.3 ms for the covariance between GF and *ϵ*_ulnar_ and between GF and *ϵ*_radial_, respectively. Some participants showed strains that lagged GF adjustments, while others showed strains that led GF adjustments. P2, the participant with the lowest mean GF, showed a strain pattern differing significantly from other participants. Indeed, P2 presented a ring of compression and dilation in *ϵ*_yy_ across oscillations leading to similar behavior in the correlation between GF and *ϵ*_radial_ and between GF and *ϵ*_ulnar_ (Fig. 6).

**Table 1.**
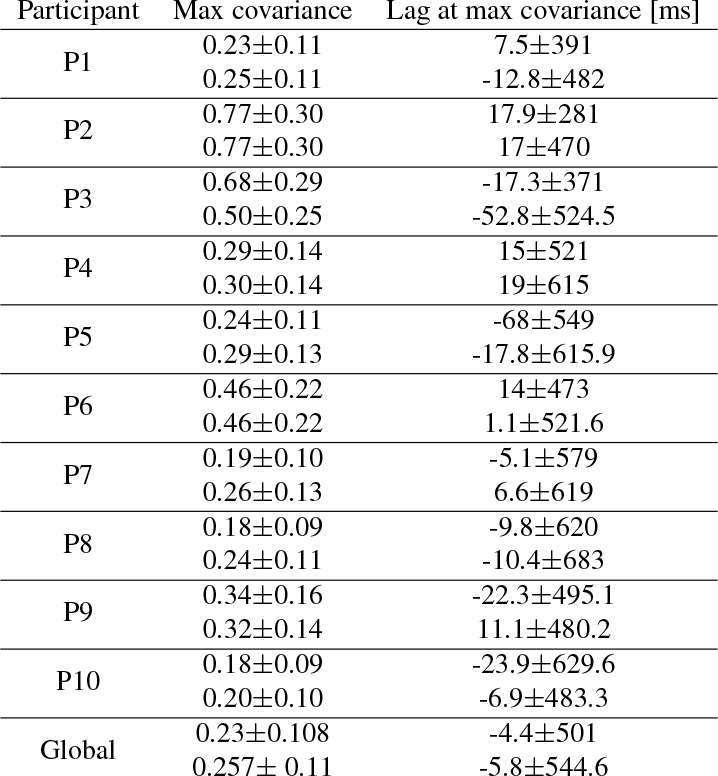
Mean *±* SD of the maximum covariance between *ϵ*_radial_ and GF, and *ϵ*_ulnar_ and GF across all trials for all participants.

**Fig. 6.**
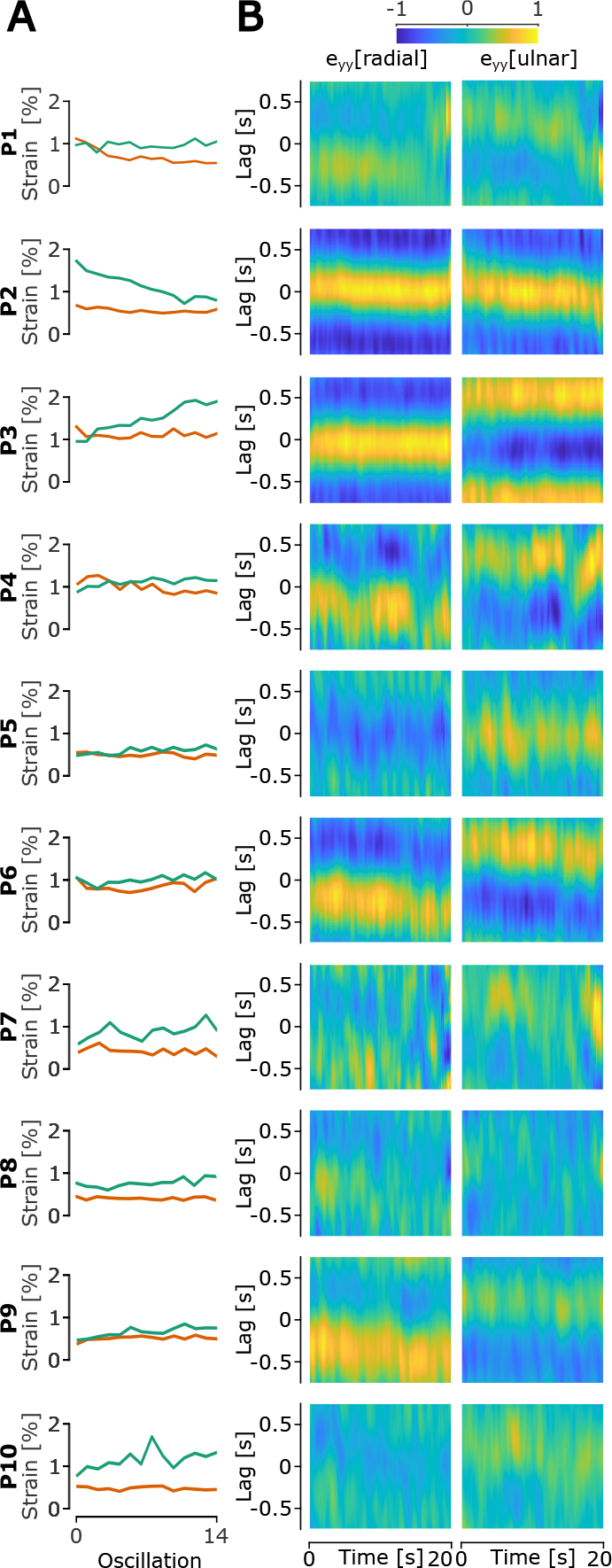
A) Mean amplitude of *ϵ*_ulnar_ (green line) and *ϵ*_radial_ (orange line) for all participants. B) Modulation of the Heatmap of *ϵ*_radial_, and *ϵ*_ulnar_ and mean crosscovariance across trials between GF and *ϵ*_radial_, and *ϵ*_ulnar_, for each participant. The cross-covariance was computed using a time window of 2.25 s. The yellow color highlights a positive covariance, and the blue color highlights a negative covariance.

We computed a similar analysis to study the covariance of peeling/laying and compression/dilation with GF, to see whether changes in the contact area size have the potential to trigger GF adjustments. Similar to what we observed in the covariance between *ϵ*_radial_, *ϵ*_ulnar_, and GF, we see that the covariance between GF and peeling/laying varies significantly across participants (Fig. 7A). Indeed, some participants showed high covariances with very small lags, while others showed small covariances with larger or highly varying lags. In terms of GF versus compression/dilation, we observed a large variation in the size of the covariance across participants, but with lags being more consistent across participants (Fig. 7B).

**Fig. 7.**
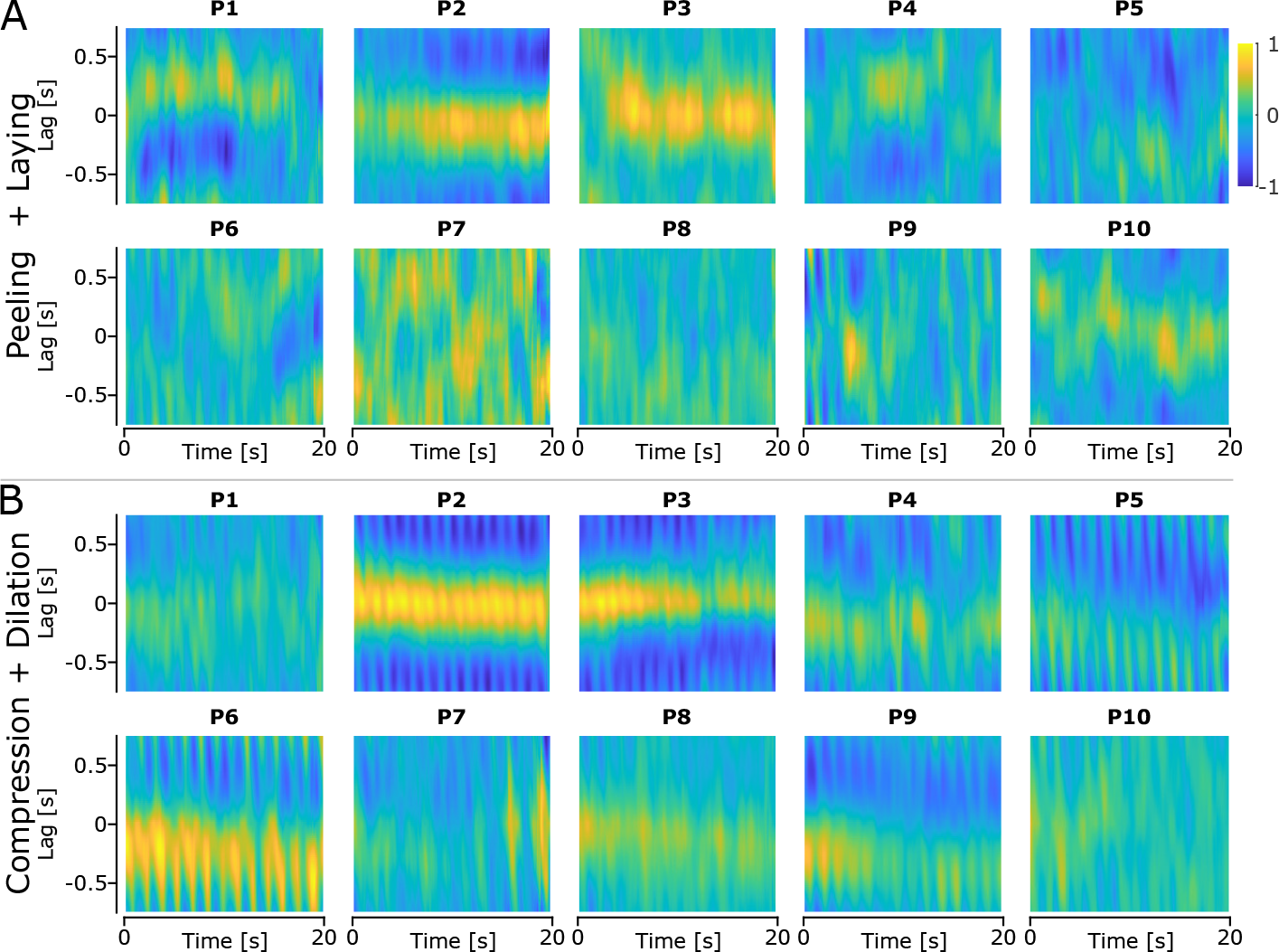
Heatmap of the mean cross-covariance between GF and the strains for each participant. The cross-covariance was computed using a time window of 2.25 s. The yellow color highlights a positive covariance, and the blue color highlights a negative covariance.

A similar cross-covariance analysis investigating the covariance between GF and LF showed that these two signals were well-correlated for all participants, with a covariance coefficient of r = 0.942 *±* 0.046 across all participants and trials. Interestingly, the lag between GF and LF was small across all participants and trials, -0.01 *±* 0.24 s (Fig. 8), highlighting how synchronous GF was in relation to LF.

**Fig. 8.**
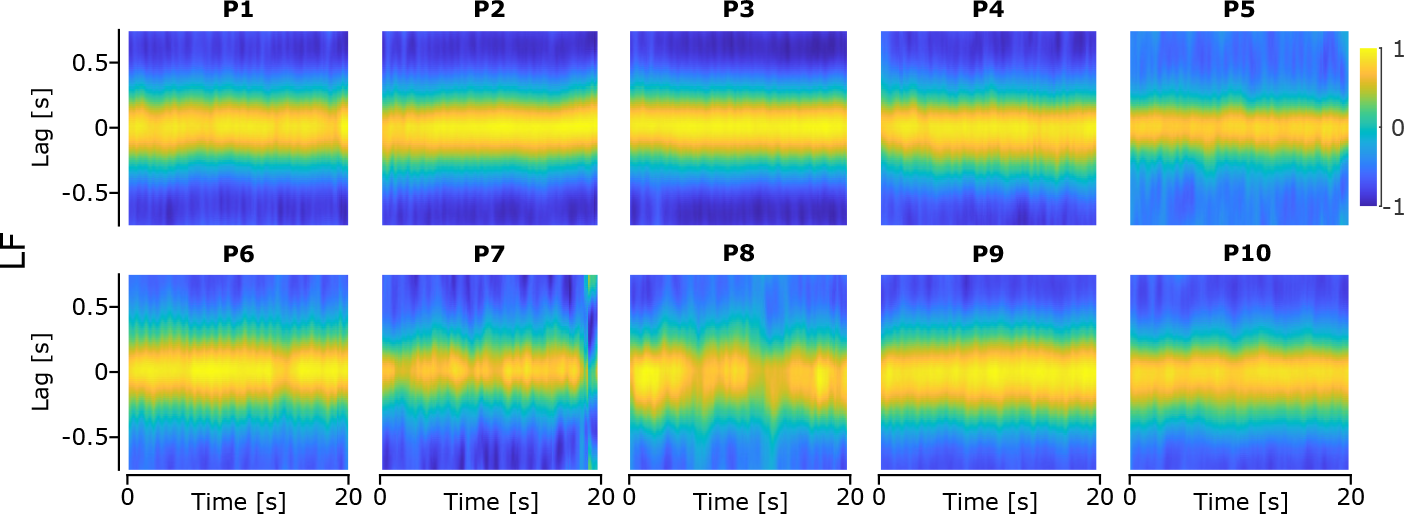
Heatmap of the mean time-windowed cross-covariance between GF and LF across all trials for each participant. The yellow color highlights a positive covariance and the blue color highlights a negative covariance.

All these results show that the behavior of the skin in the contact area, or the peeling and laying of the skin, vary significantly across participants, with all these signals not being well representative of GF adaptation for all participants. On the other hand, these results show how well synchronized GF and LF are.

Given the variability across participants, we separated them based on their mean covariance on Table 1. We set a threshold of 0.3 on this covariance and obtained two groups of participants, group 1 containing the participants with a covariance greater or equal than 0.3 and group 2 containing the participants with a covariance less than 0.3. We measured the maximum covariance between *ϵ*_*radial*_ and GF, and between *ϵ*_*ulnar*_ and GF, and extracted the lag at this maximum covariance. We observed that there was a change across trials for participants in group 1, i.e. the group with the largest covariance (Fig. 9). Given the small sample size of the groups, we did not perform statistical tests. However, the observed evolution of the max covariance in group 1 could suggest that these participants relied more on tactile feedback to adjust their GF hence increasing the relation between the two.

**Fig. 9.**
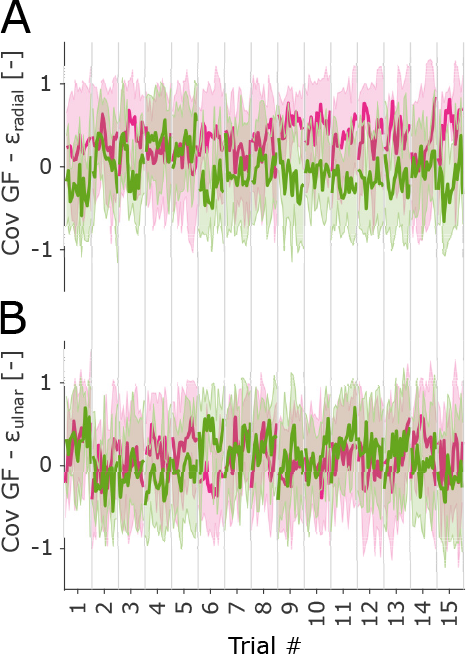
Covariance between the strains in the radial part of the finger and GF. B) Covariance between the strains in the ulnar part of the finger and GF. The pink line represents the mean and standard deviation of group 1, i.e. the group with the largest average max covariance, and the light green line represents the mean and standard deviation of group 2.

### A. Correlation between GF and strains

We extracted the correlation coefficient between GF and the strains in the radial and ulnar parts of the finger for each oscillation of each trial of every participant (Figure 2). The repeated measures ANOVA on the correlation between GF and *ϵ*_*radial*_ showed no significant main effect of either #_*osc*_ (F = 1.05, df = 13, p = 0.42) or #_*trial*_ (F = 0.88, df = 14, p = 0.58) and no significant interaction (F = 0.96, df = 182, p = 0.62). The repeated measures ANOVA on the correlation between GF and *ϵ*_*ulnar*_ showed no significant main effect of either #_*osc*_ (F = 0.45, df = 13, p = 0.94) or #_*trial*_ (F = 1.75, df = 14, p = 0.08) and no significant interaction (F = 0.87, df = 182, p = 0.86). We also performed a repeated measures ANOVA on the correlation between stains of both the radial and ulnar parts of the finger and GF of the first trial of all participants with #_*osc*_ as the within-trial variable. This ANOVA showed no significant effect of #_*osc*_ for either the correlation between *ϵ*_*radial*_ (F = 1.09, df = 14, p = 0.38) and *ϵ*_*radial*_ (F = 0.79, df = 14, p = 0.68) and GF. These results suggest that the correlation between the strains and GF did not change across oscillations or trials, indicating that participants did not change how they used strains to adjust their GF across the oscillations of a single trial, including the first trial.

## Discussion

In this paper, we present InOb, an instrumented object capable of imaging the skin of the finger pad of one finger, measuring the forces applied by the imaged finger, and measuring the accelerations and rotations experienced by the object during unconstrained manipulation. We present a use-case experiment for this object to highlight its usefulness in studying human manipulation and grip force control. In this experiment, ten participants performed vertical oscillations of 20 cm while holding InOb in a precision grip between the thumb and index finger. Participants continuously adjusted their grip force to follow the changes in the load force. We extracted various parameters quantifying the behavior of the skin in contact with the viewing plate. We observed a large variability of skin behaviors across participants, with some showing significant strains well correlated to GF adjustments while others showed small deformations or very little covariance with GF adjustments.

During the vertical oscillations, we observed that GF and LF were well synchronized with very little lag between them (Fig. 8). This indicates that participants continuously adjusted GF to match the changes in LF and maintain a safe grip with a large safety margin. Similar experiments have previously observed synchronicity between GF and LF (Delhaye et al. 2021a, Flanagan et al. 1993). This suggests that humans adjust their grip force in a predictive way so that GF changes in synchrony with LF. Recent studies have shown that the coupling between GF and LF can be context-dependent, and the strength of this coupling can change depending on the task being performed (Grover et al. 2019, 2018), with movement frequency and task difficulty influencing the GF-LF coupling. Grover *et al*. observed that more demanding tasks lead to a greater coupling between LF and GF. Indeed, movements performed at a higher frequency lead to a greater GFLF coupling than movements performed at lower frequencies. Similarly, movements performed with a larger weight lead to a greater GF-LF coupling. This suggests that the combination of weight and frequency in this experiment is demanding and requires a strong coupling between GF and LF, with very little lag between the two signals.

Previous studies have shown that tactile feedback does directly impact GF during simple grip and lift tasks (Augurelle et al. 2003, Monzée et al. 2003, Nowak et al. 2001), with GF increasing significantly when the task was performed under fingertip anesthesia. However, our data showed no consistent latency or covariance between strains, peeling/laying, compression/dilation, and GF, suggesting that the skin behavior was not the direct cause of GF adjustments observed in all trials. Moreover, when the correlation between strains and GF was computed for each individual correlation, we did not observe any particular change across trials. In some participants, we observed strain patterns during vertical oscillations (Fig. 2) that were similar to those observed in Delhaye et al. 2021a, with significant vertical strains appearing in the top and bottom parts of the finger. However, we observed significant differences in the behavior of the skin across participants, with some participants presenting large skin deformations and others presenting very small deformations. Similar variability between participant finger pad skin behavior has been observed in passive studies (du Bois de Dunilac et al. 2022, Wang and Hayward 2007). du Bois de Dunilac et al. 2022 observed that the skin deformation rate varied significantly across participants, leading to significant differences in the timing of strains during the rotational load. It is worth noting that as strains are the consequence of the contact forces, therefore strains and LF are temporally linked. However, our data suggests that this relationship is highly complex. Our data would indicate that GF control likely relies more on proprioceptive feedback or predictive control mechanisms during this task, and suggest that tactile feedback is used sparingly when the task is predictable and could have a more important role in response to unexpected perturbations, in which proprioceptive feedback is not sufficient.

To further examine the differences in skin behavior between participants, we separated our participants into two groups based on the mean covariance between GF and *ϵ*_ulnar_ and *ϵ*_radial_ they presented across all trials and all oscillations. The group with the largest covariance presented an increase in the strength of the covariance across trials, indicating that these participants might have increased their use of tactile feedback in order to adjust the GF in later oscillations and trials. However, this should be further examined in future experiments with a larger sample of participants.

However, we also observed that oscillations, where slip did occur (stick ratio < 0.5), presented a greater change in GF (greater ΔGF) than oscillations where slip did not occur. This suggests that participants did use the stick ratio and the slippage of part of the finger pad skin as feedback to adjust their grip during the manipulation, leading to greater adjustment when slips occurred. However, the oscillating pattern of the movement and of GF does not allow us to extract timing information about the timing of a change in GF following a slip of the finger pad skin.

We observed that GF decreased within and across trials as participants either adapted their grip force to the weight and inertia of the object or fatigued during the task. Grip force adaptation across trials has been observed in manipulation tasks (Giard et al. 2015, Crevecoeur et al. 2009). Despite the decrease in GF, we did not observe any significant change in strain levels within or across trials, suggesting that while participants adapted their GF, they did so in a way that did not lead to a significant reduction in grasp security.

Peeling and laying account for changes in the contact area of around 5%, which is significantly larger than the change in contact area due to compression and dilation, and average strain in the ulnar and radial parts of the fingers. We observed significant changes in peeling/laying within trials. This could indicate that the skin peeling off the contact and coming back into contact with the plate could be the main source of information about the state of the contact between the finger pad and the grasped object. Indeed, SA-I afferents are sensitive to indentation (Handler and Ginty 2021). However, the covariance and the lag at max covaraince between peeling & laying and GF varied largely across participants indicating that peeling and laying did not drive GF adjustments.

Comparing the instrumented object presented in Delhaye et al. 2021a and InOb highlights a few important differences. The device presented in Delhaye et al. 2021a weights 500 g, and the object is perfectly counterweighted to be more easily manipulable by participants, therefore having an apparent weight of 0 kg, but having the inertia of a 1 kg object. For this reason, the same experimental protocol as the one presented in this paper leads to important differences in the GF applied by participants during the experiment. Of note, we observed a distribution of mean GF that did not vary much across participants (Fig. 4B), whereas Delhaye et al. observed a much larger variation in mean GF due to the counterweighting of the object. Furthermore, due to the counterweighting mechanism, the object used by Delhaye et al. is constrained to vertical movements only, hence limiting the type of experiments that can be performed. The reduced weight of InOb (300g) and the increased mobility, allow a larger range of experiments to be performed, including experiments involving object rotations in various directions.

In order to have the capacity for unconstrained movement and limit the weight of the InOb device, a trade-off was made in terms of frame rate and pixel count when compared with the instrumented object presented in Delhaye et al. 2021a. The reduced frame rate (60 fps vs 100 fps) could limit the capacity of InOb to be used in experiments involving very fast perturbations of the object that could lead to rapid changes at the skin/object interface, for instance in the case of collisions, as skin deformations might happen too rapidly for the camera to capture. Moreover, this device is only equipped with one force/torque sensor and hence is unable to differentiate the forces applied by the thumb and index finger.

In the future, we will use InOb to perform experiments aiming to understand the role of tactile feedback during active object manipulation. We aim at using InOb in experiments requiring a task requiring reactive control of the grip force to study the influence of finger pad skin deformation on reactive adjustments of grip force to unanticipated challenges to grasp security.

## Conclusions

In this study, we introduce a new instrumented object weighing only 300 g and capable of imaging the finger pad skin, measuring the forces applied by participants on the object, and the acceleration and orientation experienced by the object. This object is made of 3D-printed elements and off-the-shelf components, allowing it to be easily manufactured. We observed strain patterns appearing on the skin during vertical oscillations but with large variability across participants. We show the use of InOb in an experiment where participants performed a vertical oscillations experiment. We show that the GF adjustments were not well correlated with strains, suggesting participants did not solely rely on tactile feedback continuously during object manipulation. InOb opens up the experimental possibilities around the study of object manipulation and could provide very useful information about how tactile feedback is used during object manipulation.

## Bibliography

Benoit Delhaye, Félicien Schiltz, Allan Barrea, Jean-Louis Thonnard, and Philippe Lefèvre. Measuring fingerpad deformation during active object manipulation. Journal of Neurophysiology, 126(4):1455–1464, October 2021a. ISSN 0022-3077, 1522-1598. doi: 10.1152/jn.00358.2021.

J.Randall Flanagan, James Tresilian, and Alan M. Wing. Coupling of grip force and load force during arm movements with grasped objects. Neuroscience Letters, 152(1-2):53–56, April 1993. ISSN 03043940. doi: 10.1016/0304-3940(93)90481-Y.

Olivier White, Noreen Dowling, R. Martyn Bracewell, and Jörn Diedrichsen. Hand Interactions in Rapid Grip Force Adjustments Are Independent of Object Dynamics. Journal of Neurophysiology, 100(5):2738–2745, November 2008. ISSN 0022-3077. doi: 10.1152/jn.90593.2008. Publisher: American Physiological Society.

Anne-Sophie Augurelle, Allan M. Smith, Thierry Lejeune, and Jean-Louis Thonnard. Importance of Cutaneous Feedback in Maintaining a Secure Grip During Manipulation of Hand-Held Objects. Journal of Neurophysiology, 89(2):665–671, February 2003. ISSN 0022-3077, 1522-1598. doi: 10.1152/jn.00249.2002.

Francis M. Grover, Patrick Nalepka, Paula L. Silva, Tamara Lorenz, and Michael A. Riley. Variable and intermittent grip force control in response to differing load force dynamics. Experimental Brain Research, 237(3):687–703, March 2019. ISSN 1432-1106. doi: 10.1007/s00221-018-5451-8. Number: 3.

Francis Grover, Maurice Lamb, Scott Bonnette, Paula L. Silva, Tamara Lorenz, and Michael A. Riley. Intermittent coupling between grip force and load force during oscillations of a hand-held object. Experimental Brain Research, 236(10):2531–2544, October 2018. ISSN 0014-4819, 1432-1106. doi: 10.1007/s00221-018-5315-2.

J. Andrew Pruszynski, Roland S. Johansson, and J. Randall Flanagan. A Rapid Tactile-Motor Reflex Automatically Guides Reaching toward Handheld Objects. Current Biology, 26(6): 788–792, March 2016. ISSN 0960-9822. doi: 10.1016/j.cub.2016.01.027.

F. Crevecoeur, A. Barrea, X. Libouton, J.-L. Thonnard, and P. Lefèvre. Multisensory components of rapid motor responses to fingertip loading. Journal of Neurophysiology, 118(1):331–343, July 2017. ISSN 0022-3077, 1522-1598. doi: 10.1152/jn.00091.2017.

Rochelle Ackerley, Ida Carlsson, Henric Wester, Håkan Olausson, and Helena Backlund Wasling. Touch perceptions across skin sites: differences between sensitivity, direction discrimination and pleasantness. Frontiers in Behavioral Neuroscience, 8, 2014. ISSN 1662-5153. doi: 10.3389/fnbeh.2014.00054.

Ingvars Birznieks, Per Jenmalm, Antony W. Goodwin, and Roland S. Johansson. Encoding of Direction of Fingertip Forces by Human Tactile Afferents. The Journal of Neuroscience, 21 (20):8222–8237, October 2001. ISSN 0270-6474, 1529-2401. doi: 10.1523/JNEUROSCI.21-20-08222.2001.

Roland S Johansson and Ingvars Birznieks. First spikes in ensembles of human tactile afferents code complex spatial fingertip events. Nature Neuroscience, 7(2):170–177, February 2004. ISSN 1097-6256, 1546-1726. doi: 10.1038/nn1177.

Giulia Corniani and Hannes P. Saal. Tactile innervation densities across the whole body. Journal of Neurophysiology, 124(4):1229–1240, October 2020. ISSN 0022-3077. doi: 10.1152/jn.00313.2020. Publisher: American Physiological Society.

Joël Monzée, Yves Lamarre, and Allan M. Smith. The Effects of Digital Anesthesia on Force Control Using a Precision Grip. Journal of Neurophysiology, 89(2):672–683, February 2003. ISSN 0022-3077. doi: 10.1152/jn.00434.2001. Publisher: American Physiological Society.

T. André, V. Lévesque, V. Hayward, P. Lefèvre, and J.-L. Thonnard. Effect of skin hydration on the dynamics of fingertip gripping contact. Journal of The Royal Society Interface, 8(64): 1574–1583, November 2011. doi: 10.1098/rsif.2011.0086. Publisher: Royal Society.

Benoit Delhaye, Philippe Lefèvre, and Jean-Louis Thonnard. Dynamics of fingertip contact during the onset of tangential slip. Journal of The Royal Society Interface, 11(100):20140698, November 2014. ISSN 1742-5689, 1742-5662. doi: 10.1098/rsif.2014.0698.

M. Tada and T. Kanade. An imaging system of incipient slip for modelling how human perceives slip of a fingertip. In The 26th Annual International Conference of the IEEE Engineering in Medicine and Biology Society, volume 1, pages 2045–2048, September 2004. doi: 10.1109/IEMBS.2004.1403601.

Vincent Levesque and Vincent Hayward. Experimental Evidence of Lateral Skin Strain During Tactile Exploration. 2003.

Benoit Delhaye, Allan Barrea, Benoni B. Edin, Philippe Lefèvre, and Jean-Louis Thonnard. Surface strain measurements of fingertip skin under shearing. Journal of The Royal Society Interface, 13(115):20150874, February 2016. ISSN 1742-5689, 1742-5662. doi: 10.1098/rsif.2015.0874.

Sophie du Bois du Bois de Dunilac, David Córdova Bulens, Philippe Lefèvre, Stephen J. Redmond, and Benoit P. Delhaye. Biomechanics of Finger Pad Response under Torsion. preprint, Neuroscience, November 2022.

Benoit Delhaye, Ewa Jarocka, Allan Barrea, Jean-Louis Thonnard, Benoni Edin, and Philippe Lefèvre. High-resolution imaging of skin deformation shows that afferents from human fingertips signal slip onset. eLife, 10:e64679, April 2021b. ISSN 2050-084X. doi: 10.7554/eLife.64679. Publisher: eLife Sciences Publications, Ltd.

Felicien Schiltz, Benoit P Delhaye, Jean-Louis Thonnard, and Philippe Lefevre. Grip Force is adjusted at a level that maintains an upper bound on partial slip across friction conditions during object manipulation. IEEE Transactions on Haptics, pages 1–1, 2021. ISSN 2329-4051. doi: 10.1109/TOH.2021.3137969. Conference Name: IEEE Transactions on Haptics.

Serena Bochereau, Brygida Dzidek, Michael Adams, and Vincent Hayward. Characterizing and Imaging Gross and Real Finger Contacts under Dynamic Loading. IEEE Transactions on Haptics, 10(4):456–465, October 2017. ISSN 1939-1412, 2329-4051, 2334-0134. doi: 10.1109/TOH.2017.2686849.

Laurence Willemet, Nicolas Huloux, and Michaël Wiertlewski. Efficient tactile encoding of object slippage. Scientific Reports, 12(1):13192, August 2022. ISSN 2045-2322. doi: 10.1038/s41598-022-16938-1. Number: 1 Publisher: Nature Publishing Group.

Heba Khamis, Hafiz Malik Naqash Afzal, Jennifer Sanchez, Richard Vickery, Michaël Wiertlewski, Stephen J. Redmond, and Ingvars Birznieks. Friction sensing mechanisms for perception and motor control: passive touch without sliding may not provide perceivable frictional information. Journal of Neurophysiology, 125(3):809–823, March 2021. ISSN 0022-3077. doi: 10.1152/jn.00504.2020. Publisher: American Physiological Society.

Allan Barrea, David Cordova Bulens, Philippe Lefevre, and Jean-Louis Thonnard. Simple and Reliable Method to Estimate the Fingertip Static Coefficient of Friction in Precision Grip. IEEE Transactions on Haptics, 9(4):492–498, October 2016. ISSN 1939-1412. doi: 10.1109/TOH.2016.2609921.

Bruce D Lucas and Takeo Kanade. ITERATIVE IMAGE REGISTRATION TECHNIQUE WITH AN APPLICATION TO STEREO VISION. volume 2, pages 674–679, 1981.

Jianbo Shi and Carlo Tomasi. Good features to track. In Proceedings of the IEEE Computer Society Conference on Computer Vision and Pattern Recognition, pages 593–600, 1994. ISBN 0818658274. doi: 10.1109/cvpr.1994.323794.

Jean yves Bouguet. Pyramidal implementation of the lucas kanade feature tracker. Intel Corporation, Microprocessor Research Labs, 2000.

Dennis A. Nowak, Joachim Hermsdörfer, Stefan Glasauer, Jens Philipp, Ludger Meyer, and Norbert Mai. The effects of digital anaesthesia on predictive grip force adjustments during vertical movements of a grasped object. European Journal of Neuroscience, 14 (4):756–762, 2001. ISSN 1460-9568. doi: 10.1046/j.0953-816x.2001.01697.x. _eprint: https://onlinelibrary.wiley.com/doi/pdf/10.1046/j.0953-816x.2001.01697.x.

Qi Wang and Vincent Hayward. In vivo biomechanics of the fingerpad skin under local tangential traction. Journal of Biomechanics, 40(4):851–860, January 2007. ISSN 0021-9290. doi: 10.1016/j.jbiomech.2006.03.004.

T. Giard, F. Crevecoeur, J. McIntyre, J.-L. Thonnard, and P. Lefèvre. Inertial torque during reaching directly impacts grip-force adaptation to weightless objects. Experimental Brain Research, 233(11):3323–3332, November 2015. ISSN 1432-1106. doi: 10.1007/s00221-015-4400-z.

F. Crevecoeur, J. L. Thonnard, and P. Lefèvre. Forward models of inertial loads in weightlessness. Neuroscience, 161(2):589–598, June 2009. ISSN 0306-4522. doi: 10.1016/j.neuroscience.2009.03.025.

Annie Handler and David D. Ginty. The mechanosensory neurons of touch and their mechanisms of activation. Nature Reviews Neuroscience, 22(9):521–537, September 2021. ISSN 1471-0048. doi: 10.1038/s41583-021-00489-x. Number: 9 Publisher: Nature Publishing Group.

